# scPADGRN: A preconditioned ADMM approach for reconstructing dynamic gene regulatory network using single-cell RNA sequencing data

**DOI:** 10.1101/799189

**Authors:** Xiao Zheng, Yuan Huang, Xiufen Zou

## Abstract

Disease development and cell differentiation both involve dynamic changes; therefore, the reconstruction of dynamic gene regulatory networks (DGRNs) is an important but difficult problem in systems biology. With recent technical advances in single-cell RNA sequencing (scRNA-seq), large volumes of scRNA-seq data are being obtained for various processes. However, most current methods of inferring DGRNs from bulk samples may not be suitable for scRNA-seq data. In this work, we present scPADGRN, a novel DGRN inference method using time-series scRNA-seq data. scPADGRN combines the preconditioned alternating direction method of multipliers with cell clustering for DGRN reconstruction. It exhibits advantages in accuracy, robustness and fast convergence. Moreover, a quantitative index called Differentiation Genes’ Interaction Enrichment (DGIE) is presented to quantify the interaction enrichment of genes related to differentiation. From the DGIE scores of relevant subnetworks, we infer that the functions of embryonic stem (ES) cells are most active initially and may gradually fade over time. The communication strength of known contributing genes that facilitate cell differentiation increases from ES cells to terminally differentiated cells. We also identify several genes responsible for the changes in the DGIE scores occurring during cell differentiation based on three real single-cell datasets. Our results demonstrate that single-cell analyses based on network inference coupled with quantitative computations can reveal key transcriptional regulators involved in cell differentiation and disease development.

**Author summary:** Single-cell RNA sequencing (scRNA-seq) data are gaining popularity for providing access to cell-level measurements. Currently, time-series scRNA-seq data allow researchers to study dynamic changes during biological processes. This work proposes a novel method, scPADGRN, for application to time-series scRNA-seq data to construct dynamic gene regulatory networks, which are informative for investigating dynamic changes during disease development and cell differentiation. The proposed method shows satisfactory performance on both simulated data and three real datasets concerning cell differentiation. To quantify network dynamics, we present a quantitative index, DGIE, to measure the degree of activity of a certain set of genes in a regulatory network. Quantitative computations based on dynamic networks identify key regulators in cell differentiation and reveal the activity states of the identified regulators. Specifically, Bhlhe40, Msx2, Foxa2 and Dnmt3l might be important regulatory genes involved in differentiation from mouse ES cells to primitive endoderm (PrE) cells. For differentiation from mouse embryonic fibroblast cells to myocytes, Scx, Fos and Tcf12 are suggested to be key regulators. Sox5, Meis2, Hoxb3, Tcf7l1 and Plagl1 critically contribute during differentiation from human ES cells to definitive endoderm cells. These results may guide further theoretical and experimental efforts to understand cell differentiation processes and explore cell heterogeneity.

## Introduction

In systems biology, the reconstruction of dynamic gene regulatory networks (DGRNs) has proven to be a crucial tool for understanding processes related to disease development and cell differentiation, such as hematopoietic specification [1], T cell activation [2], influenza infection, acute lung injury, and type 2 diabetes [3]. DGRNs specify links between genes over time. By exploring the differences in dynamic networks, researchers are able to comprehend the mechanisms causing complex diseases [3], etc.

Recently, large quantities of single-cell RNA sequencing (scRNA-seq) data have been obtained for various biological processes due to advances in sequencing techniques [4–7]. However, most current methods of inferring DGRNs for bulk samples may not be suitable for scRNA-seq data. For example, methods involving ordinary differential equations (ODEs) become invalid since the biological meaning of a sample changes from the average for several cells in bulk data to the value for a single cell. Several individual cells can be sequenced at once, causing the form of the gene expression data to change from a single vector to several vectors, or a matrix. The cells sequenced at different time points are different. It is not possible to describe the dynamics of a single cell because that cell does not even exist at the next time point. However, the dynamics of cells at the cluster level can be described by ODEs. This is a compromise approach to exploring cell heterogeneity information based on single-cell data.

In this work, we present scPADGRN, a novel method of inferring DGRNs from time-series scRNA-seq data. scPADGRN combines the preconditioned alternating direction method of multipliers (PADMM) with cell clustering for DGRN reconstruction. The cell clustering process includes ranking cells in accordance with their pseudotimes and merging cells into clusters. Our optimization model considers network precision, network sparsity and network continuity. The PADMM is used to solve the optimization model to obtain the DGRN. Multiple matrices are updated, and three subproblems are solved by the PADMM algorithm in each iteration.

Simulated data and three real datasets concerning cell differentiation have been used to test the performance of scPADGRN. We propose a quantity called Differentiation Genes’ Interaction Enrichment (DGIE) to quantify the changes in the interactions of a certain set of genes in a DGRN. First, we chose genes involved in the same biological processes or KEGG pathways to visualize subnetworks of DGRNs and computed their DGIE scores. Then, we selected all genes known to contribute to the process of cell differentiation and computed the corresponding DGIE scores. We also identified several genes responsible for the drastic changes in the DGIE scores in each dataset. These genes might be key regulators in cell differentiation. Our results demonstrate that single-cell analyses based on network inference coupled with quantitative computations can reveal key transcriptional regulators in cell differentiation and disease development.

## Materials and methods

### Simulated datasets

In this section, we describe the simulation of cluster-specific data *Y* = [*Y* (1), …, *Y* (*N*)]. First, we simulated the *X*_1_ values in time-series single-cell data *X* = [*X*_1_, …, *X*_*N*_] using the scRNA-seq simulation tool Splatter [12]. After setting appropriate numbers of genes (*m*), cells (*n*) and cell clusters (*r*), we generated the initial gene expression data *X*_1_ using Splatter. Then, we constructed cluster-specific data *Y* (1) by merging vectors (cells) belonging to the same cluster into a single vector, representing the gene expression value of the cluster. The next step was to generate the *Y* (*t*), 2 ≤ *t* ≤ *N*. We defined the dynamic network {*A*(1), …, *A*(*N* − 1)} in the form of random 0-1 matrices and *Y* (*t* + 1) = *A*(*t*)*Y* (*t*) + *Y* (*t*), 1 ≤ *t* ≤ *N* − 1. After these steps, cluster-specific data *Y* = [*Y* (1), …, *Y* (*N*)] were obtained.

In the experiments on the simulated data, there were two main questions of concern: how noise and the number of clusters *r* affect the network accuracy. To answer these two questions, we conducted two separate experiments. In the first experiment, we set the number of genes to 100, 200, 300, 400 and 500, individually. The number of cells was set to 10 times the number of genes, and the number of clusters was equal to the number of genes. We also set the number of time points to *N* = 5. Thus, we obtained corresponding cluster-specific data *Y* = [*Y* (1),…, *Y* (*N*)]. Here, we considered the noise to be independent and to follow a Gaussian distribution with a mean value of *µ* = 0 and a standard deviation of *σ* = 0.01, 0.02 or 0.05. By adding noise to the cluster-specific data *Y* = [*Y* (1), …, *Y* (*N*)], we obtained noisy cluster-specific datasets.

In the second experiment, we set the number of genes to 200 and 400. The number of cells was set to 10 times the number of genes, and the number of clusters was varied from 40 to 200 and from 40 to 400, separately. The number of time points was again set to *N* = 5.

### Three real datasets

Three time-series scRNA-seq datasets concerning cell differentiation were obtained from [8], with pseudotimes inferred by Monocle [9]. Dataset 1 was derived from mouse embryonic stem cells (ES cells) differentiating to primitive endoderm (PrE) cells [5]. A total of 356 cells, which were sequenced at 0, 12, 24, 48 and 72 h, were used. Dataset 2 was derived from mouse embryonic fibroblast cells differentiating to myocytes [6]. A total of 405 cells were sequenced at days 0, 2, 5, and 22. Dataset 3 contains data from 758 cells sequenced at 0, 12, 24, 36, 72 and 96 h. Dataset 3 was derived from human ES cells differentiating to definitive endoderm cells [7]. For all three real data examples, reference networks from the Transcription Factor Regulatory Network database (http://www.regulatorynetworks.org) were used to validate the inferred networks.

### DGRN reconstruction

In this work, we propose a novel DGRN inference method called scPADGRN. The framework of scPADGRN is shown in Fig 1. Two main steps are needed to infer DGRNs. First, we cluster scRNA-seq data for different cells based on cell pseudotrajectories to convert single-cell-level data into cluster-level data. Details on the cell clustering process are provided in Fig 2. Second, the PADMM method is used to solve the optimization problem with the reshaped data. Fig 3 shows a flowchart of the PADMM algorithm.

**Fig 1.**
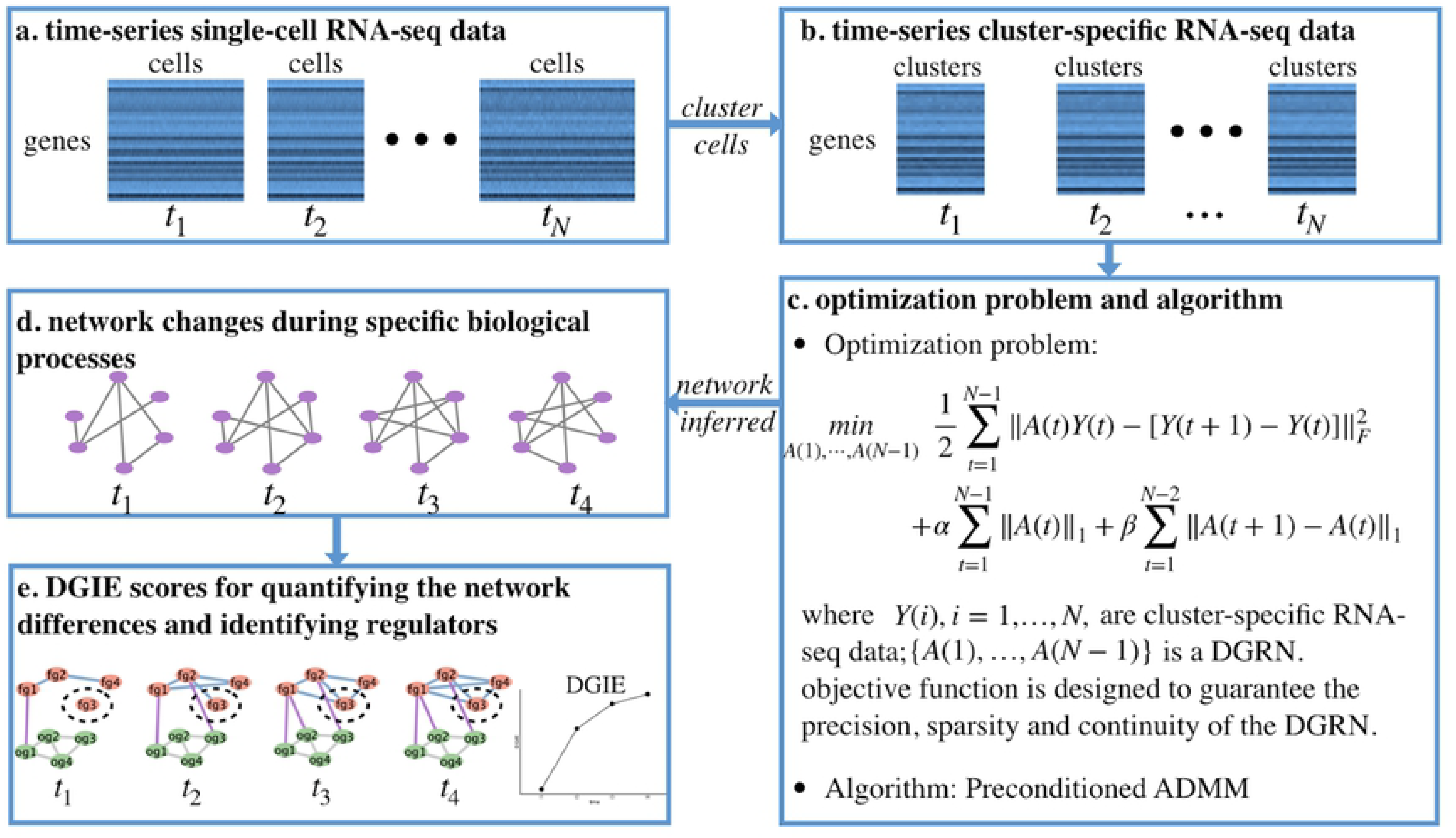
Framework of scPADGRN. (a) Time-series scRNA-seq data. Several cells are sequenced at each time point. (b) Time-series cluster-specific RNA-seq data. The same clusters exist at each time point. (c) Optimization problem and algorithm. Three features of DGRNs are considered in the optimization problem: precision, sparsity and continuity. The PADMM method is used to solve the optimization problem. (d) Network changes during specific biological processes. The purple nodes represent the genes involved in the same biological processes. Several links change during a given process. (e) DGIE scores for quantifying the network differences and identifying regulators. The nodes shown in pink are functional genes (fg). The nodes shown in green are other genes (og). The DGIE score measures the activity state of the functional genes. The blue and purple links are used to compute the DGIE scores. In this toy model, the DGIE score increases over time since the interactions of the functional genes become more intense. The circled gene, fg3, is the identified key transcriptional regulator.

**Fig 2.**
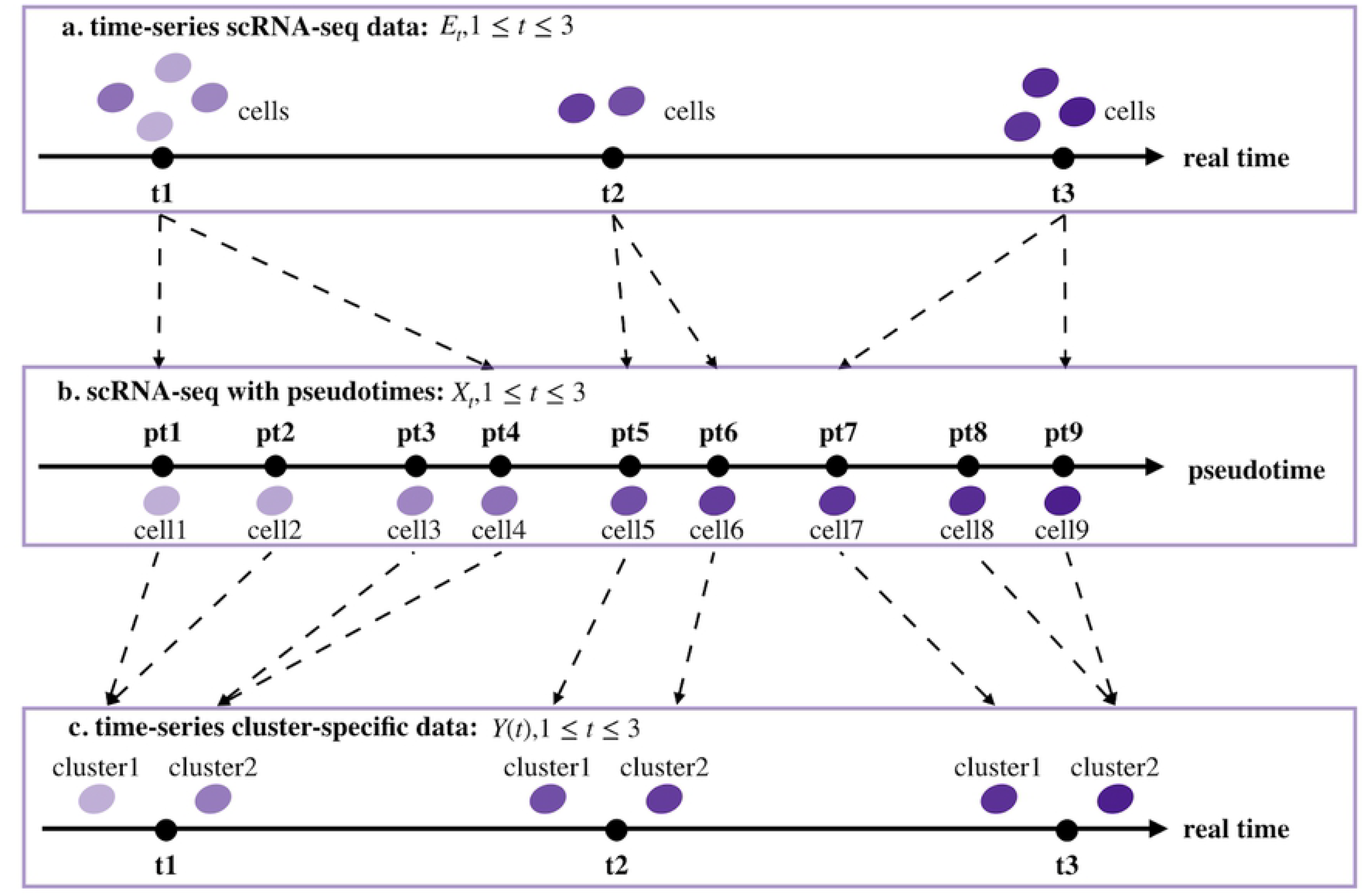
Clustering process for data conversion. (a) Time-series scRNA-seq data *E*_*t*_, 1 ≤ *t* ≤ 3. Several cells are sequenced at each time point. (b) Corresponding scRNA-seq data *X*_*t*_, 1 ≤ *t* ≤ 3, under pseudotimes. The cells are arranged on a pseudotime line. (c) Time-series cluster-specific data. The same clusters exist at each time point on the real timeline.

**Fig 3.**
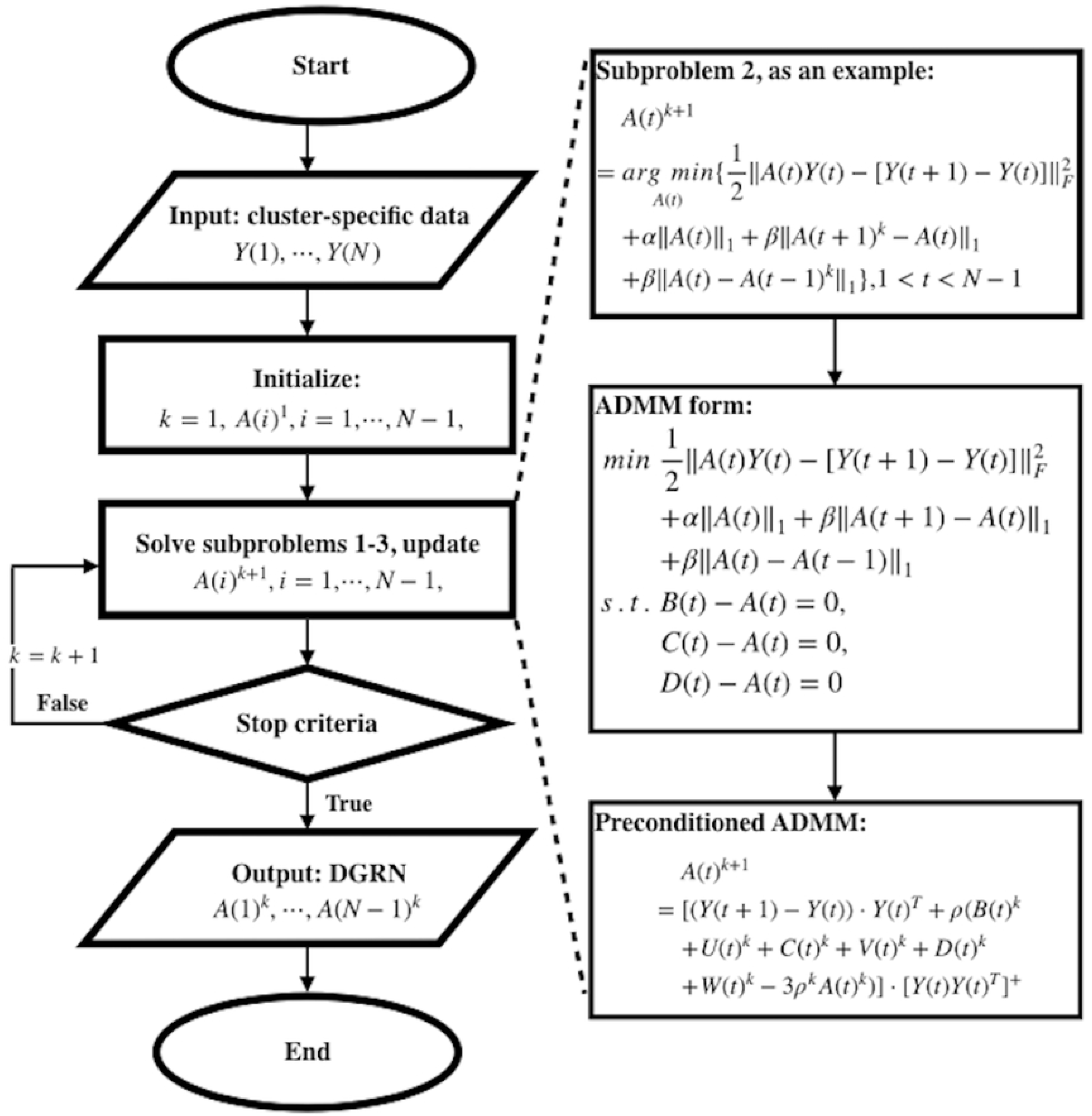
Flowchart of the PADMM method. The processes include inputting the cluster-specific data *Y* (1), …, *Y* (*N*), initializing the variables, and updating the *A*(*t*), 1 ≤ *t* ≤ *N* − 1. The PADMM algorithm is used to solve all three subproblems in each iteration.

#### Data conversion: from single-cell-level data to cluster-level data

First, we introduce the time-series scRNA-seq data. The time-series scRNA-seq data are denoted by *E*_*t*_, 1 ≤ *t* ≤ *N*, representing matrices of gene expression values at *N* different time points. The *E*_*t*_, 1 ≤ *t* ≤ *N*, are *m*_*t*_ × *n*_*t*_ numerical matrices whose rows represent the genes (features) and whose columns represent the cells (samples) at time *t*. Element (*E*_*t*_)_*ij*_ of *E*_*t*_ is the expression value of the *i*-th gene in the *j*-th cell at time *t*. Generally, the genes at each time point are identical. Namely, their features are identical, and the number of features is *m*_1_ = *m*_2_ = … = *m*_*N*_ = *m*. In contrast, the cells at each time point are totally different individuals. Usually, the number of samples *n*_*i*_ is not equal to *n*_*j*_ if *i* ≠ *j*.

In Fig 2, an example with three time points is used to illustrate the two steps of data conversion. The first step is to acquire the pseudotrajectory information of all cells and rank the cells at each real time point from early to late stages in accordance with their pseudotimes. Namely, we realign the columns of *E*_*t*_, 1 ≤ *t* ≤ *N*. The reshaped data are denoted by *X*_*t*_, 1 ≤ *t* ≤ *N*. Mature technologies such as Monocle [9] can be employed to infer the cell pseudotrajectories. As part of this step, we project the cells on the real timeline to cells on a pseudotime line.

The second step is to cluster the cells on the pseudotime line into clusters on the real timeline. In detail, the conversion process includes the following operations. We set the number of clusters *r* equal to the minimum of the numbers of cells *n*_*t*_, 1 ≤ *t* ≤ *N*. For the realigned *X*_*t*_, we compute the distance between the gene expression vectors of every pair of adjacent cells. Then, we take the largest *n*_*t*_ − *r* distances among the obtained *n*_*t*_ − 1 distances and link their corresponding cells. We consider linked cells to belong to the same cluster. In this way, *r* ordered clusters are obtained. For the *r* ordered clusters of *X*_*t*_, we use *y*_*j*_(*t*) to denote the gene expression of the *j*-th cluster at time *t*. *y*_*j*_(*t*) is a column vector consisting of the row means of the matrix composed of the cells in the *j*-th cluster at time *t*.

We adopt the notation *Y* (*t*) = [*y*_1_(*t*), …, *y*_*k*_(*t*)], 1 ≤ *t* ≤ *N*, where the *Y* (*t*), 1 ≤ *t* ≤ *N*, are *m* × *r* matrices representing the gene expression levels of the *r* clusters at time *t*. Through these steps, we convert the time-series single-cell data *X* = [*X*_1_, …, *X*_*N*_] into time-series cluster-specific gene expression data *Y* = [*Y* (1),…, *Y*(*N*)].

Since the cells at each time point are different, it is difficult to describe the expression dynamics at the single-cell level. For example, suppose that cell 1 is sequenced at *t*_1_ and cell 2 is sequenced at *t*_2_, where *t*_1_ < *t*_2_. Cell 1 will be destroyed upon being sequenced at *t*_1_. Therefore, cell 1 does not correspond to any cells at *t*_2_. One feasible solution is to describe the dynamics at the cluster level; in this way, little information about cell heterogeneity is lost.

#### Optimization of DGRN

The expression dynamics of the *i*-th gene can be described by the following ODE:

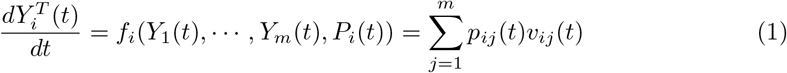

where *Y*_*i*_(*t*) is a continuous vector in time *t*, representing the *i*-th row of *Y* (*t*). *Y*_*i*_(*t*) represents the expression level of the *i*-th gene. *v*_*ij*_(*t*) and *p*_*ij*_(*t*) denote the reaction and the reaction rate, respectively, from the *j*-th gene to the *i*-th gene at time *t*. *P*_*i*_(*t*) is a parameter set.

To construct the DGRN, we need to search for the optimal parameter set 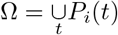 in Eq (1). This problem can be converted into the problem of finding a set Ω to fit the simulation results to the experimental results. We consider the augmentation of cluster-specific data between two adjacent time points. Let the sequencing times be denoted by *t*_*r*_, and let *t*_*r*_ = 1, 2, …, *N*. The optimization problem is as follows:

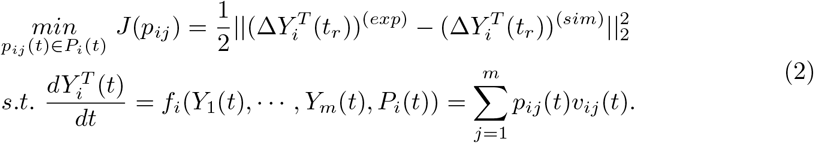

The objective of problem (2) is to optimize the augmentation of the gene expression of the *i*-th gene at time *t*_*r*_, and it is a nonlinear dynamic optimization problem (DOP), which is one of the most difficult types of optimization problems to solve. To simplify this problem, we presume that the interactions among genes between two adjacent discrete time points *t*_*r*_ and *t*_*r*_ + 1 are linear. We use a piecewise linearization technique to approximate Eq (1):

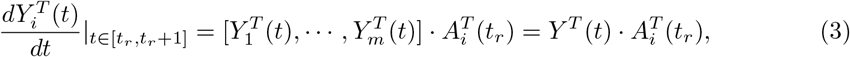

where *A*_*i*_(*t*_*r*_) is the *i*-th row of the *m* × *m* matrix *A*(*t*_*r*_). Thus, the optimization problem (2) is converted into

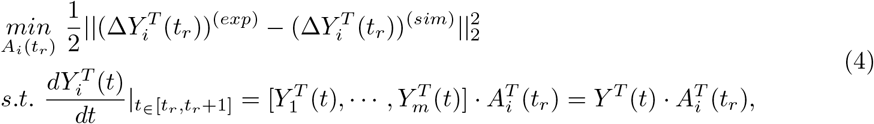

where 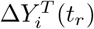 is the difference in gene expression between *t*_*r*_ and *t*_*r*_ + 1. (⋅)^(*exp*)^ and (⋅)^(*sim*)^ denote the experimental and simulated results, respectively.

The objective of problem (4) is to optimize the parameters of the dynamics of the *i*-th gene at time *t*_*r*_. In the next step, we sum all *m* genes and all *N* time points simultaneously.

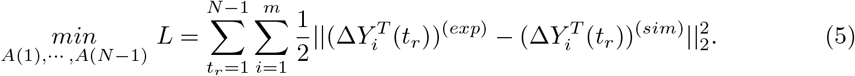

With Eq (3), we also have the following approximation:

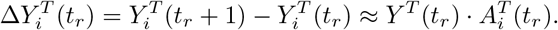

Then, the objective function *L* in problem (5) can be written as

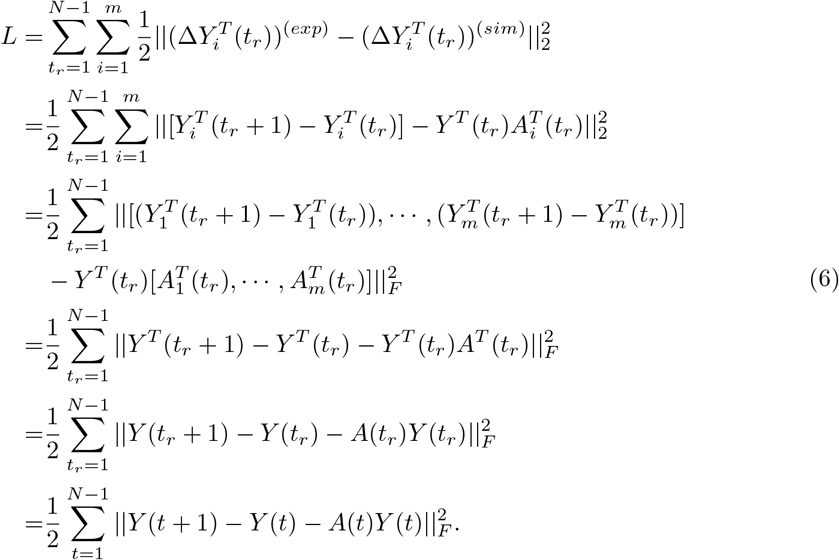

In the DGRN {*A*(1), …, *A*(*N* − 1)}, the nodes stand for genes, and the links stand for gene regulatory relationships between genes. The DGRN is a directed dynamic network whose positive and negative links correspond to activation and suppression relationships, respectively. Usually, DGRNs are sparse and continuous. In other words, most parameters in problem (5) will be zero, and the differences between the network states at two adjacent time points should be slight. Therefore, we define the following optimization problem:

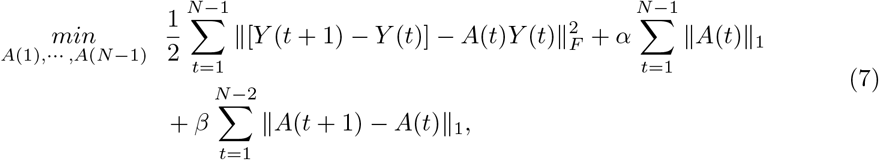

where the first term evaluates the precision of problem (5), the second term is the *L*_1_-norm of the dynamic network to guarantee the sparsity of the network, and the third term imposes the continuity assumption on the dynamic network states at consecutive time points. Both sparsity and continuity need to be considered in biological networks [3]. The parameters *α* and *β* are tuning parameters that control the penalties for sparsity and continuity, respectively.

### PADMM Algorithm

There are *N* − 1 matrices that need to be optimized in problem (7). We use the alternating descent method to iteratively solve the problem. In each iteration, we update the *N* − 1 matrices sequentially. For each matrix *A*(*t*), 1 ≤ *t* ≤ *N* − 1, we update *A*(*t*) while keeping the other *N* − 2 matrices fixed.

In the *k*-th iteration, for the update of *A*(*t*), 1 ≤ *t* ≤ *N*, there are three different cases, each corresponding to a different subproblem.

- Subproblem 1 When *t* = 1, there are three terms in the objective function.

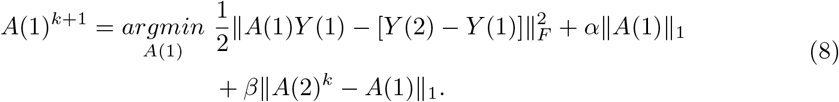
- Subproblem 2 When *t* = 2, …, *N* − 2, there are four terms in the objective function.

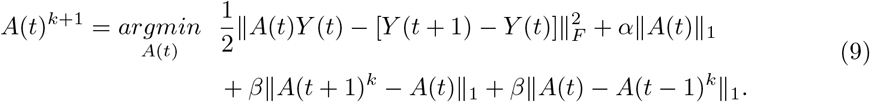
- Subproblem 3 When *t* = *N* − 1, there are three terms in the objective function.

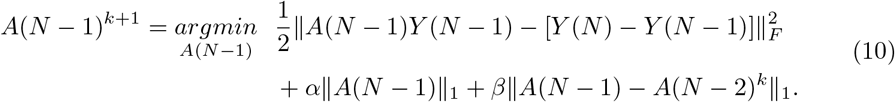

The PADMM is a variation of the alternating direction method of multipliers (ADMM, [11]). Before introducing the PADMM, we present the ADMM algorithm for solving these three subproblems. The scaled ADMM [11] is employed here since it is a more convenient form.

For subproblem 1, first, we convert it into ADMM form:

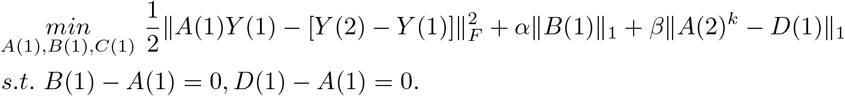

Its augmented Lagrangian is

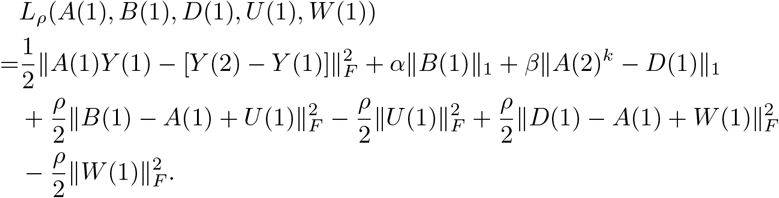

The iterations are as follows:

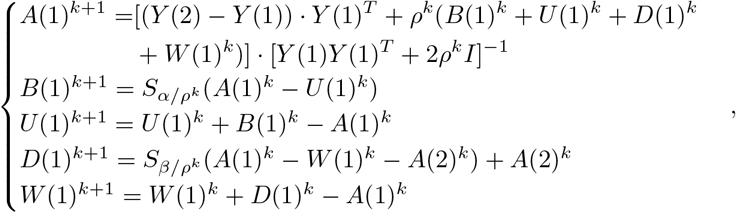

where the soft thresholding operator *S* is defined as

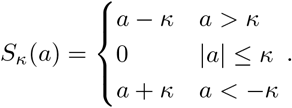

For subproblem 2, we convert it into ADMM form:

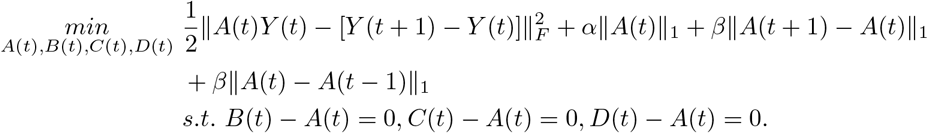

Its augmented Lagrangian is

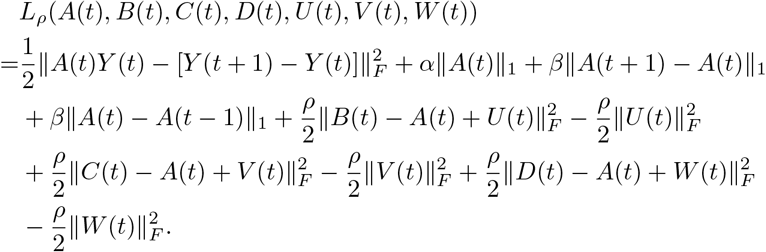

The iterations are as follows:

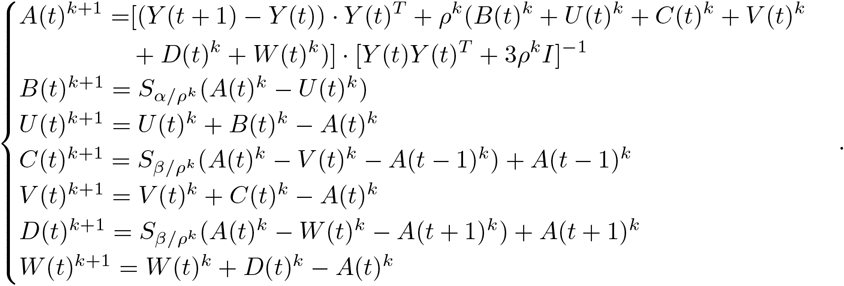

Subproblem 3 is similar to subproblem 1. When *t* = *N* − 1,

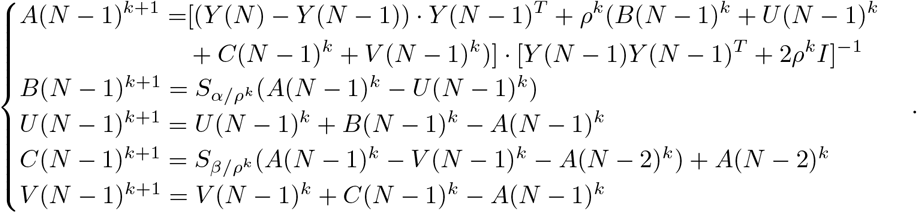

With some adjustments to the ADMM described above, one can use the PADMM to achieve a faster computation speed. Proper preconditioning processes are applied for the computation of *A*(*t*), 1 ≤ *t* ≤ *N* − 1.

The form of the iterations of *A*(*t*), 1 ≤ *t* ≤ *N* − 1, arises from 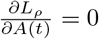. Consider *t* = 1 (subproblem 1) as an example.

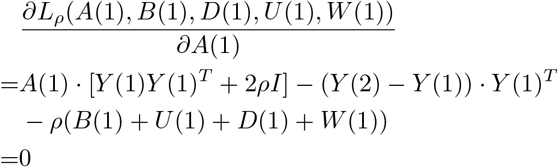

is equivalent to

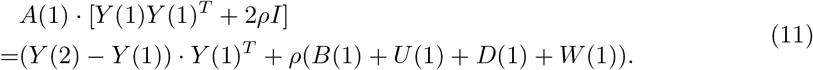

Usually, in the ADMM, the update *A*(1) of takes the form

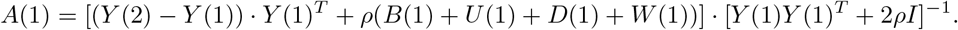

With the proposed preconditioning,, we add −2*ρA*(1) to both sides of Eq (11). As the result,

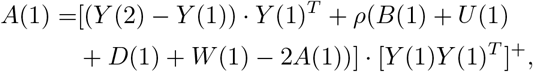

where (*M*)^+^ denotes the general inverse of the matrix *M*, in case *M* is singular.

Similarly, we can obtain the PADMM iterations of *A*(*t*), 1 ≤ *t* ≤ *N* − 1 for all subproblems as follows:

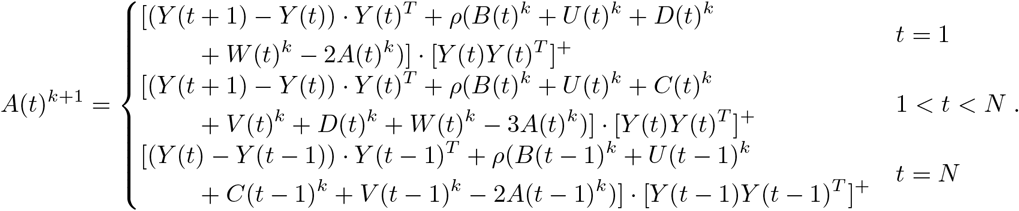

The [*Y*(*t*)*Y* (*t*)^*T*^]^+^, 1 ≤ *t* ≤ *N*, are unchanged in all iterations; therefore, they can be stored as constants. Hence, the PADMM can save *N* matrix inversion computations in every iteration except the first. Singular value decomposition is used to compute the general inverses of the [*Y* (*t*)*Y* (*t*)^*T*^]^+^, 1 ≤ *t* ≤ *N*. Proper preconditioning makes the computation of the matrix inverses easier while maintaining an equivalent precision. Details on the theoretical results can be found in [10].

#### Parameter selection

- Algorithm parameters The number of clusters *r* is set to the minimum among the numbers of cells at all time points. When *t* = 1, we take *A*(*t*), *U* (*t*), *V* (*t*) and *W* (*t*) as zero matrices and *B*(*t*), *C*(*t*) and *D*(*t*) as random matrices. A maximum number of iterations *M* and a relative error threshold *ϵ* are set. Iteration is terminated when the maximum number of iterations *M* is reached or when 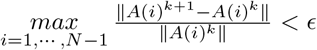. The parameter *ρ* is chosen such that *ρ*^*k*+1^ = *ρ*^*k*^/2. For details on the algorithm parameters, please refer to [11].
- Model selection The chosen model parameters *α* and *β* strongly affect the network structure. Bayesian information criterion (BIC) can be used to optimize the parameters *α* and *β* [3]. Let *L*^∗^ denote the objective function of optimization problem (7). We formulate the BIC optimization problem as follows:

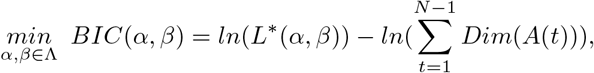

where Λ = {*α*_0_, …, *α*_*l*_}. Here, *α*_*i*+1_ = *α*_*i*_*ρ*, *i* = 0, …, *l* − 1, with 0 < *ρ* < 1. *Dim*(⋅) denotes the dimensionality of the argument in parentheses, and we consider this quantity to take non-negative values, as follows: *Dim*(*A*(*t*)) = …, where *δ* > 0 is a threshold.
- Choice of network thresholds Once the weighted adjacent matrices are computed, different network thresholds may lead to different network structures. We assume that the first network state of the dynamic network has the same average degree as the reference network, whose links have been confirmed by biological experiments.

### Analysis of network differences

#### DGIE scores for measuring changes in the interactions of a certain set of genes in a DGRN

To quantify the differences in the dynamic network states over time, we propose the DGIE score. Suppose that we want to study the progress of cell differentiation. Let the DGRN states be denoted by *G*_*t*_ = (*V*_*t*_, *E*_*t*_), 1 ≤ *t* ≤ *N* − 1, where *N* − 1 is the number of network states. Suppose that the vertex set is 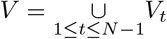. We divide the vertex set *V* into two disjoint subsets *V*_(1)_ and *V*_(2)_. *V*_(1)_ is the set of genes that are known to contribute to processes related to cell differentiation, including cell growth, proliferation, and development. This information is available in gene annotation databases, such as Metascape [14]. Another possible choice for *V*_(1)_ is to select genes that belong to the same pathway. In this case, the DGIE scores can help identify the activation states of this pathway. After *V*_(1)_ is determined, *V*_(2)_ is the set of the remaining genes.

We define the DGIE score as

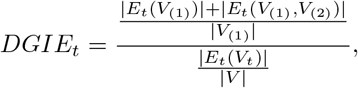

where 1 ≤ *t* ≤ *N* − 1 and *DGIE*_*t*_ is an *N* − 1-dimensional array. *E*_*t*_(*V*_(1)_) is the edge set of the subgraph whose vertex set is *V*_(1)_ in the *t*-th network state of the DGRN. *E*_*t*_(*V*_(1)_, *V*_(2)_) is the edge set of the bigraph whose vertex sets are *V*_(1)_ and *V*_(2)_ in the *t*-th network state of the DGRN. *| ⋅ |* is the number of elements of a set. The denominator 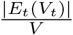 in the definition of *DGIE*_*t*_ is the ratio of the number of links in *G*_*t*_ to the number of genes in *V*, and it is used to alleviate the effects caused by different numbers of links at different time points. The numerator 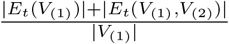 in the definition of *DGIE*_*t*_ is the ratio of the sum of the number of links in *V*_(1)_ and the number of links between *V*_(1)_ and *V*_(2)_ to the number of genes in *V*_(1)_. The definition of *DGIE*_*t*_ mainly concerns the sum of the number of links in *V*_(1)_ and the number of links between *V*_(1)_ and *V*_(2)_. To minimize the effects of parameters such as |*V*_(1)_|, |*E*_*t*_(*V*_(1))_| and *|V|*, we define *DGIE*_*t*_ as shown above to measure the communication ability of the genes in *V*_(1)_.

#### Local differences: dynamic subnetworks and DGIE scores for specific biological processes

Extracting subnetworks from a DGRN is an efficient way to clearly see the network differences. We choose genes related to the same biological process or pathway and extract their corresponding subnetworks. By comparing these subnetworks with the reference network, one can easily see the corresponding network differences from the subnetworks themselves, including changes in interactions and directions.

Then, we can compute the DGIE scores of the subnetworks and look for invariant characteristics. For real data applications, we focus here on subnetworks related to ES cell differentiation processes.

#### Global differences: DGIE scores of all known contributing genes

For the biological processes described by DGRNs, for example, differentiation from mouse ES cells to PrE cells, many genes contribute to related tasks, such as the regulation of embryonic development, the determination of cell fate, cell cycle regulation, and the encoding of de novo DNA methyltransferases. Information about gene annotation can help to identify these known contributing genes. With these contributing genes as *V*_1_, computing the DGIE scores enables us to learn more about changes in the communication strength of these genes.

#### Identifying key regulators responsible for changes in DGIE scores

To investigate the mechanisms underlying drastic changes in DGIE scores, it is important to identify the genes which are responsible for those changes. By removing one gene from *V*_(1)_ at a time, we can observe the resulting changes in the DGIE scores. If the removed gene is irrelevant to the changes in the DGIE scores, the DGIE scores should still drastically vary. On the other hand, if the DGIE scores are almost identical at each time point after the removal of a certain gene, then this gene should be considered responsible for the originally observed variations. Furthermore, with the removal of a combination of genes (a complex), the standard deviation of the DGIE scores at all time points may also be reduced to a rather low level. In this case, the removed complex is our target. The method of complex identification involves the following steps. First, the differentiation-related genes are ranked in accordance with their ability to reduce the standard deviation of the DGIE scores. Then, the first *d, d* = 1, 2, …, genes in the ranked list are taken as a complex, and the DGIE scores after the removal of this complex are calculated. This process is repeated until the standard deviation of the DGIE scores no longer decreases. The corresponding complex is what we are looking for.

After identifying the complex responsible for the changes in the DGIE scores for each dataset, we can then investigate the role of complexes in DGRNs. We extract links adjacent to these genes at each time point and draw the corresponding differential network. By comparing the differential network with the reference network, some of the links can be confirmed to be biologically meaningful. The links without such confirmation are the links that we predict to be crucial to the biological process.

## Results

In this section, we report simulation experiments carried out to demonstrate the effectiveness of the proposed algorithms. Then, we infer and analyze DGRNs based on three real scRNA-seq datasets related to cell differentiation processes.

### Numerical experiments on simulated data

#### Effects of noise level on network accuracy

The methods used to construct the simulated data are described in the materials and methods section. Here, two algorithms, the ADMM and PADMM algorithms, were tested. The runtime, numbers of iterations, reconstruction errors, and areas under the receiver operating characteristic curves (AUCs) were calculated. Table 1 shows the results for 300 and 500 genes. The complete results are listed in S1 Table.

**Table 1.** Effects of noise level on network accuracy.

| **gene number=300** |  |  |  |  |  |  |  |  |  |  |  |
| --- | --- | --- | --- | --- | --- | --- | --- | --- | --- | --- | --- |
| Noise | Method | Time(s) | #iteration | Reconstruction error |  |  |  | AUC |  |  |  |
|  |  |  |  | t1 | t2 | t3 | t4 | t1 | t2 | t3 | t4 |
| 0 | ADMM | 24.676 | 24 | 2.000 | 3.420 | 4.815 | 6.270 | 0.998 | 1.000 | 1.000 | 1.000 |
|  | PADMM | 9.854 | 32 | 2.000 | 3.420 | 4.815 | 6.270 | 0.998 | 1.000 | 1.000 | 1.000 |
| 0.01 | ADMM | 23.896 | 25 | 8.595 | 16.626 | 18.136 | 23.125 | 0.998 | 1.000 | 1.000 | 1.000 |
|  | PADMM | 9.006 | 30 | 8.608 | 16.744 | 18.235 | 23.220 | 0.998 | 1.000 | 1.000 | 1.000 |
| 0.02 | ADMM | 23.952 | 25 | 16.402 | 37.052 | 41.270 | 38.369 | 0.998 | 1.000 | 1.000 | 1.000 |
|  | PADMM | 8.511 | 28 | 16.422 | 37.328 | 41.522 | 38.461 | 0.998 | 1.000 | 1.000 | 1.000 |
| 0.05 | ADMM | 23.925 | 25 | 53.181 | 81.860 | 69.184 | 71.881 | 0.994 | 0.985 | 0.992 | 0.990 |
|  | PADMM | 8.550 | 28 | 53.430 | 82.395 | 69.383 | 72.058 | 0.994 | 0.984 | 0.992 | 0.990 |
| **gene number=500** |  |  |  |  |  |  |  |  |  |  |  |
| Noise | Method | Time(s) | #iteration | Reconstruction error |  |  |  | AUC |  |  |  |
|  |  |  |  | t1 | t2 | t3 | t4 | t1 | t2 | t3 | t4 |
| 0 | ADMM | 134.612 | 34 | 3.486 | 4.881 | 7.242 | 9.316 | 0.999 | 1.000 | 1.000 | 1.000 |
|  | PADMM | 34.831 | 32 | 3.486 | 4.881 | 7.242 | 9.316 | 0.999 | 1.000 | 1.000 | 1.000 |
| 0.01 | ADMM | 156.080 | 34 | 9.877 | 25.193 | 33.505 | 41.704 | 0.999 | 1.000 | 1.000 | 1.000 |
|  | PADMM | 37.173 | 30 | 9.877 | 25.187 | 33.494 | 41.693 | 0.999 | 1.000 | 1.000 | 1.000 |
| 0.02 | ADMM | 138.125 | 34 | 37.224 | 45.211 | 68.268 | 83.807 | 0.998 | 1.000 | 1.000 | 0.999 |
|  | PADMM | 35.510 | 30 | 37.212 | 45.196 | 68.250 | 83.788 | 0.998 | 1.000 | 1.000 | 0.999 |
| 0.05 | ADMM | 105.719 | 26 | 64.804 | 70.041 | 130.689 | 158.855 | 0.995 | 1.000 | 0.988 | 0.972 |
|  | PADMM | 35.457 | 28 | 64.871 | 70.079 | 130.933 | 158.895 | 0.995 | 1.000 | 0.988 | 0.972 |

Runtime shows that PADMM is faster than ADMM by 60.07%-76.12%. Reconstruction errors suggest that PADMM and ADMM share the similar precision. AUC measures accuracy from the perspective of binary classification, and PADMM and ADMM both perform well on AUC.

From the results in Table 1 and S1 Table, reconstruction errors increase and AUCs decrease as the noise level increases, as expected. There is little difference on AUC for ADMM and PADMM while PADMM reduces runtime by 67.77% on average. From the perspective of binary classification, these two algorithms are both capable of identifying most links.

#### Effects of the number of cell clusters on network accuracy

We used two simulation datasets to examine the effects of the number of cell clusters. The number of clusters *r* is crucial because a smaller *r* corresponds to a smaller number of known variables. More specifically, the ratio of the number of known variables to the number of unknown variables is 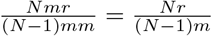 in problem (7). We need to know the extent of the effect of the number of clusters.

The runtime, numbers of iterations, reconstruction errors and AUCs were computed. Table 2 shows the results obtained for 200 genes with numbers of clusters ranging from 40 to 200. The complete results are listed in S2 Table.

**Table 2.** Effects of cell cluster numbers on network accuracy.

| gene number=200 |  |  |  |  |  |  |  |  |  |  |  |
| --- | --- | --- | --- | --- | --- | --- | --- | --- | --- | --- | --- |
| Cluster number | Method | Time(s) | #iteration | Reconstruction error |  |  |  | AUC |  |  |  |
|  |  |  |  | t1 | t2 | t3 | t4 | t1 | t2 | t3 | t4 |
| 200 | ADMM | 8.921 | 25 | 1.533 | 1.828 | 2.996 | 4.172 | 1.000 | 1.000 | 1.000 | 1.000 |
|  | PADMM | 3.886 | 33 | 1.533 | 1.828 | 2.996 | 4.171 | 1.000 | 1.000 | 1.000 | 1.000 |
| 160 | ADMM | 5.819 | 17 | 8.990 | 8.627 | 8.852 | 9.023 | 1.000 | 1.000 | 1.000 | 1.000 |
|  | PADMM | 2.236 | 18 | 8.990 | 8.627 | 8.852 | 9.023 | 1.000 | 1.000 | 1.000 | 1.000 |
| 120 | ADMM | 5.080 | 17 | 12.748 | 12.541 | 12.573 | 12.523 | 1.000 | 1.000 | 1.000 | 1.000 |
|  | PADMM | 1.272 | 12 | 12.749 | 12.542 | 12.573 | 12.523 | 1.000 | 1.000 | 1.000 | 1.000 |
| 80 | ADMM | 5.317 | 17 | 15.460 | 15.614 | 15.329 | 15.408 | 0.995 | 0.998 | 0.997 | 0.999 |
|  | PADMM | 1.476 | 10 | 15.460 | 15.614 | 15.329 | 15.408 | 0.995 | 0.998 | 0.997 | 0.999 |
| 40 | ADMM | 4.982 | 17 | 17.658 | 17.777 | 17.812 | 17.784 | 0.970 | 0.979 | 0.984 | 0.984 |
|  | PADMM | 1.509 | 10 | 17.658 | 17.777 | 17.812 | 17.784 | 0.970 | 0.979 | 0.984 | 0.984 |

PADMM and PADMM can identify DGRNs accurately when the number of clusters are far less than the number of genes. PADMM is faster than ADMM by 66.99% on average.

As seen from the results in Table 2 and S2 Table, reconstruction errors increase and AUCs decrease with a decreasing number of clusters. When the number of clusters decreases to 2/5 of the number of genes (Table 2), the AUC remains above 0.99, which is sufficiently high. When the number of clusters decreases to 1/10 of the number of genes (with 400 genes), the AUC remains above 0.92, as shown in S2 Table. These results show that both algorithms are able to identify most of the links in a DGRN with a rather small number of clusters. The ADMM and PADMM algorithms both maintain good precision, as shown in the simulation experiments. In addition, PADMM is faster than ADMM by an average of 66.99%, as seen in Table 2.

As seen from the results of both simulation experiments, the ADMM and PADMM are both able to identify links in dynamic networks despite the occurrence of noise and a small number of clusters. However, the PADMM is superior to the ADMM in terms of runtime. Therefore, for the real data analyses reported below, we used the PADMM.

### Applications to real scRNA-seq data

#### Dataset 1: mouse ES cells to PrE cells

In accordance with the described methods for inferring DGRNs, we obtained the DGRN for dataset 1, as shown in S1 Fig. Furthermore, we visualized subnetworks of genes involved in GO:0048863 stem cell differentiation. We selected genes that are involved in both the reference network and the DGRN. Subnetworks with eight genes are shown in Fig 4.

**Fig 4.**
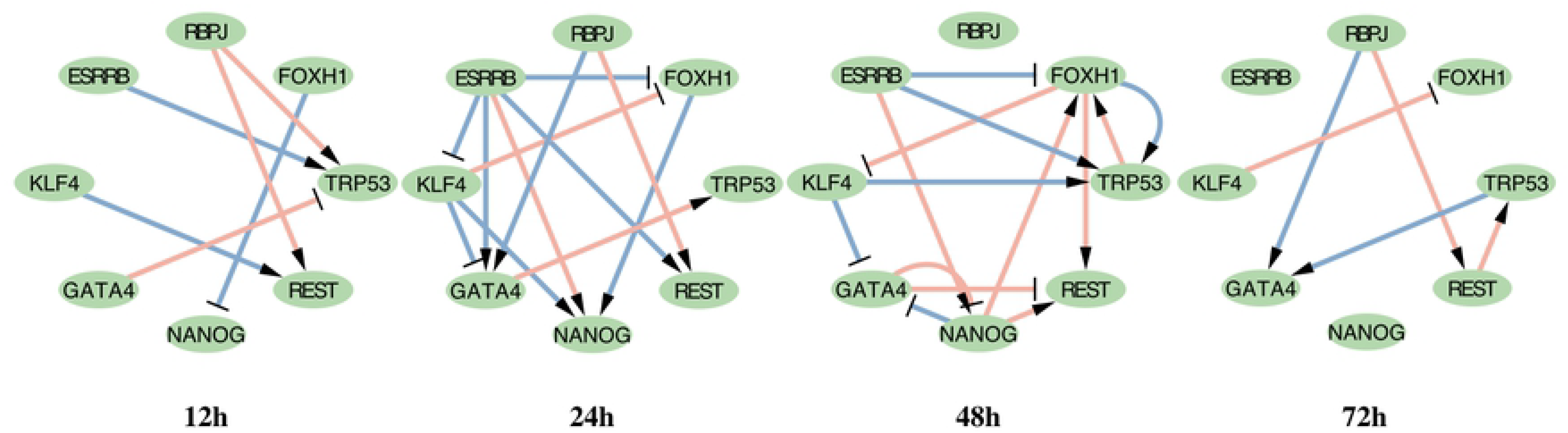
Dataset 1: Subnetworks of DGRNs with genes in GO:0048863 stem cell differentiation. Gene nodes are genes in GO:0048863. Pink links are TF-TF interactions confirmed by biological experiments. Links with arrow and ‘T’ are positive and negative interactions, respectively.

All network figures presented in this work were plotted using Cytoscape [13]. Transcription factor (TF)-TF interactions confirmed by biological experiments are marked in pink. Links marked with arrows and ‘T’ symbols represent positive and negative interactions, respectively. In these subnetworks, RBPJ and ESRRB regulate the other six genes without being regulated themselves. TRP53 and REST are activated at all times. FOXH1 is suppressed beginning at 24 h. GATA4 is both activated and suppressed beginning at 24 h.

The DGIE scores of the genes in Fig 4 are shown in Fig 7(a). Datasets 1 and 3 describe differentiation processes for mouse and human ES cells, respectively. Therefore, we chose GO:0048863 stem cell differentiation for dataset 1 and hsa04550 signaling pathways regulating the pluripotency of stem cells for dataset 3, among other biological processes and KEGG pathways that are less relevant to the differentiation of ES cells. By observing the DGIE scores of genes in subnetworks, we may learn the activation states of the corresponding biological processes and KEGG pathways.

Fig 7(a) and Fig 7(b) both show a decreasing tendency. As seen from the definition of DGIE, the DGIE score measures the communication ability of a certain set of genes. The observed decrease in the DGIE score indicates a decrease in the communication ability of the cells involved in GO:0048863 stem cell differentiation. In other words, this biological process becomes less activated over time. This result is consistent with the biological phenomenon if we hypothesize that the differentiation of ES cells influences the communication ability of related genes and vice versa. According to textbooks on cell biology, once ES cells begin to differentiate, they are no longer ES cells. The degree of differentiation of the cells becomes higher at that time. Therefore, it is natural to assume that the communication ability of these genes begins to fade since the cells become increasingly dissimilar to ES cells as time goes by. The same decreasing tendency is observed in both mouse and human ES cell differentiation, as shown in Fig 7.

Next, we consider the process of the differentiation of mouse ES cells to PrE cells. We take all known contributing genes as *V*_1_. The DGIE scores are shown in Fig 8 (a). The observed increasing tendency suggests that the interactions within the genes in *V*_1_ and between *V*_1_ and *V*_2_ intensify over time.

In fact, the differentiation from ES cells to PrE cells is only an early stage of the differentiation of stem cells into terminally differentiated cells. Similar increasing tendencies are also observed in datasets 2 and 3. From the increasing tendency in Fig 8, we can infer that functions that facilitate cell differentiation, including cell growth, proliferation, and development, are gradually turned on. The DGIE score is a tool for determining the activation states of functions at the molecular level.

S4 Fig shows boxplots of the DGIE scores when a gene or complex is removed from *V*_1_. We identify four genes, BHLHE40, MSX2, FOXA2 and DNMT3L, as targets.

According to the gene annotation information available from the Metascape database [14], BHLHE40 is involved in the control of the circadian rhythm and cell differentiation. MSX2 may promote cell growth under certain conditions. DNMT3L is crucial for embryonic development. Similar family members of FOXA2 regulate metabolism and play a role in the differentiation of pancreas and liver cells in mice. It is known that endoderm cells will differentiate into pancreas and liver cells. Thus, it is also natural to infer that FOXA2 may play a key role in early ES cell differentiation even before pancreas and liver cells are formed.

In addition, let 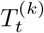 denote the set of genes with the top *k* largest degrees in the DGRN at time *t*, with *k* = 10 and 50. We compare 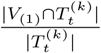 with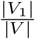. The results are shown in S4 Table. In S4 Table(A), it is clear that differentiation-related genes are denser among top-degree nodes, and top-degree nodes are usually regarded as possessing higher influence in a complex network.

S7 Fig shows the differential network formed based on the union of links that appear between the complex and other genes only once from 12 h to 72 h. Counts of the confirmed links in the differential network are shown in S5 Table. The unconfirmed links may play important roles in the biological process.

#### Dataset 2: mouse embryonic fibroblast cells to myocytes

For dataset 2, we visualized subnetworks of genes involved in GO:0061614 pri-miRNA transcription by RNA. mi-RNA is hypothesized to regulate approximately one-third of human genes; therefore, we are interested in how genes interact with others to facilitate pri-miRNA transcription by RNA. Nine genes were selected, as shown in Fig 5.

**Fig 5.**
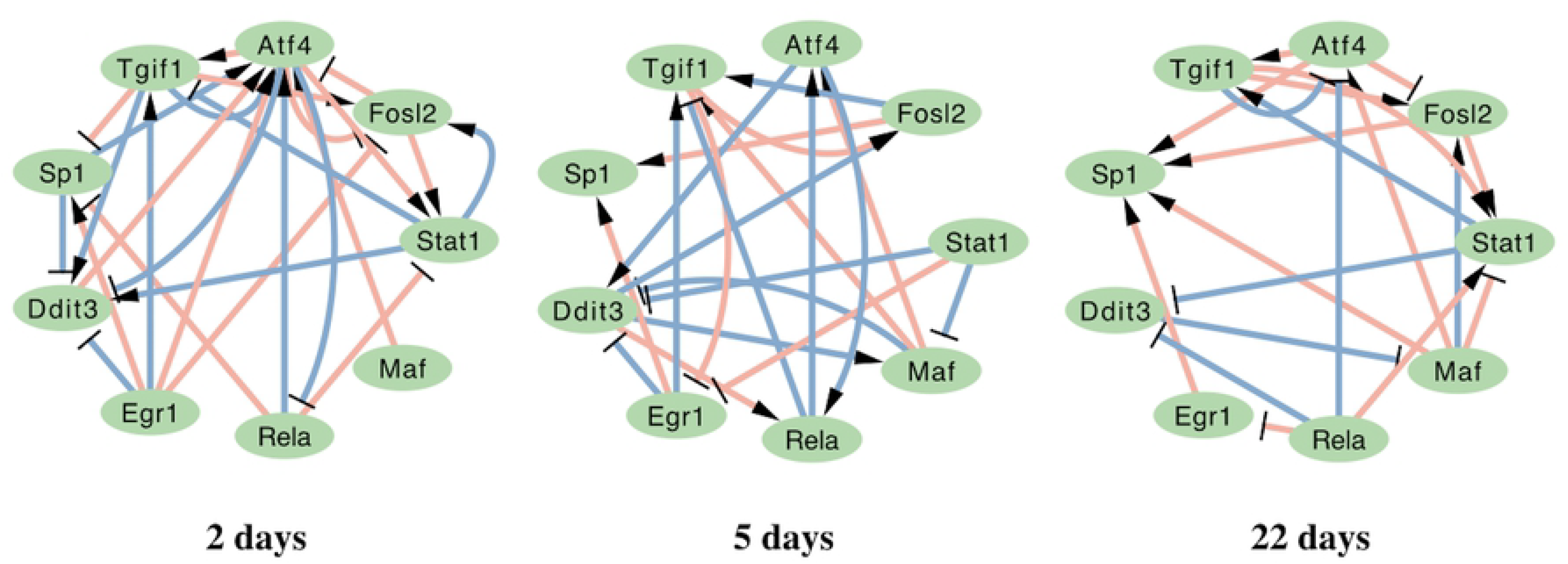
Dataset 2: Subnetworks of DGRNs with genes in GO:0061614 pri-miRNA transcription by RNA. Gene nodes are genes in GO:0061614. Pink links are TF-TF interactions confirmed by biological experiments. Links with arrow and ‘T’ are positive and negative interactions, respectively.

In these subnetworks, ATF4, TGIF1, SP1, DDIT3 and FOSL2 are activated and suppressed at all times. EGR1 is suppressed beginning at 5 days. MAF is both suppressed and activated beginning at 5 days. A full image of the DGRN states is shown in S2 Fig.

The DGIE scores of all known contributing genes are shown in Fig 8(b). As in the case of dataset 1, we perceive an increasing tendency of the DGIE scores over time. It is worth mentioning that dataset 2 does not describe cell differentiation from ES cells directly. Instead, it describes cell differentiation from less differentiated cells to myocytes, which are terminally differentiated cells.

For the process of differentiation from ES cells to terminally differentiated cells, we know that the DGIE scores increase from the ES cells to more highly differentiated cells, such as the PrE cells in dataset 1. The DGIE scores also increase from less differentiated cells (fibroblasts) to terminally differentiated cells (myocytes). Thus, it would not be too bold to infer that the communication strength of the known contributing genes increases from ES cells to terminally differentiated cells. Although no biological experiments yet confirm this claim, we present this speculation from the perspective of dynamic network analysis.

S5 Fig shows boxplots of the DGIE scores when a gene or complex is removed from *V*_(1)_. We identify three genes as key transcriptional regulators: Scx, Fos and Tcf12. According to the gene annotation information available from the Metascape database, Scx regulates collagen type I gene expression in cardiac fibroblasts and myofibroblasts. Fos proteins regulate cell proliferation, differentiation, and transformation. Tcf12 is expressed in many tissues, including skeletal muscle.

#### Dataset 3: human ES cells to definitive endoderm cells

Dataset 3 describes differentiation from human ES cells to definitive endoderm cells. As in the case of dataset 1, we focused on biological processes or KEGG pathways that are directly involved in stem cell differentiation. Therefore, we chose ten genes in hsa04550 signaling pathways regulating the pluripotency of stem cells for visualization. The subnetworks are shown in Fig 6.

**Fig 6.**
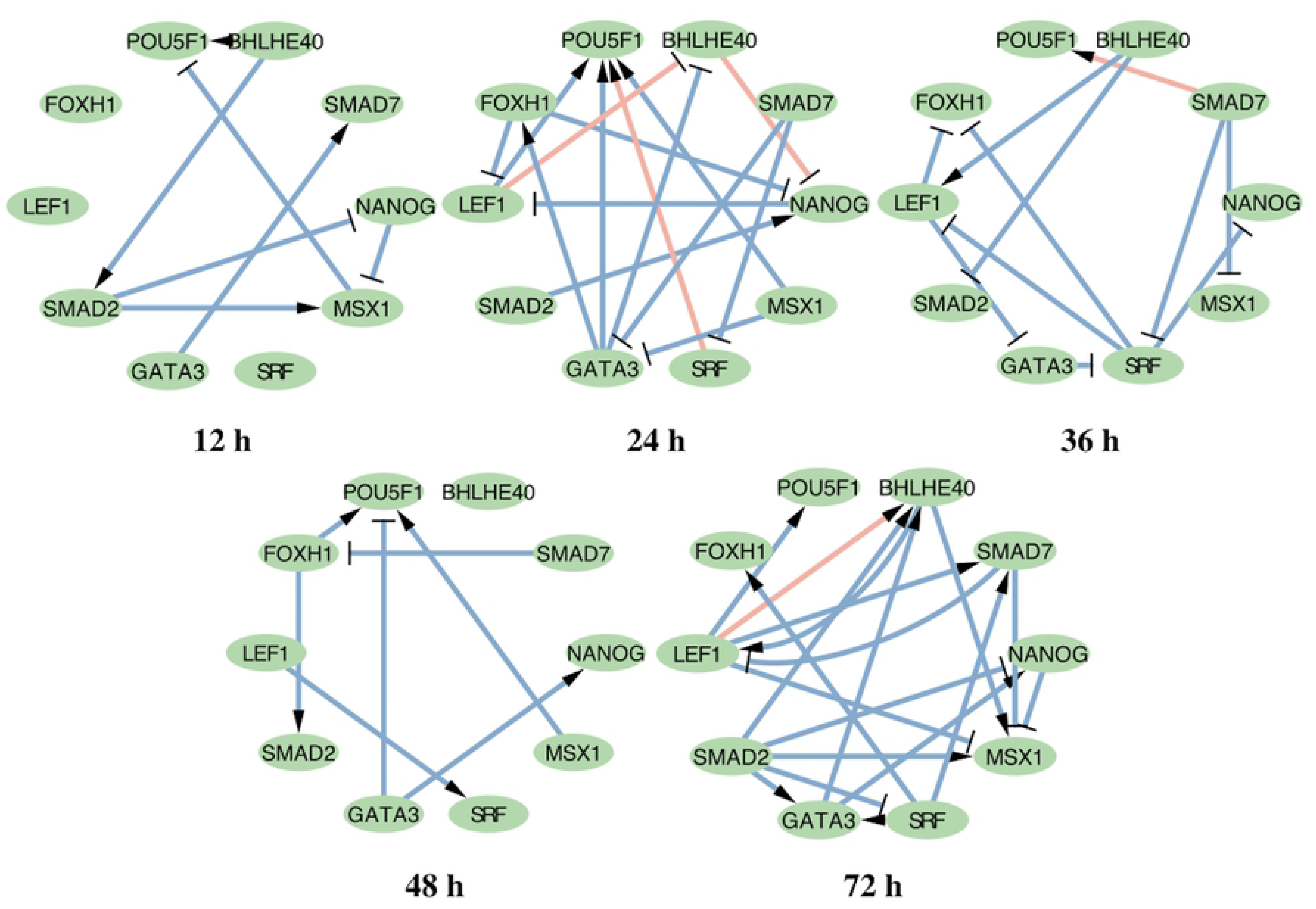
Dataset 3: Subnetworks of DGRNs with genes in hsa04550 signaling pathways regulating pluripotency of stem cells. Gene nods are genes in hsa04550 signaling pathways regulating pluripotency of stem cells. Pink links are TF-TF interactions confirmed by biological experiments. Links with arrow and ‘T’ are positive and negative interactions, respectively.

The subnetworks in Fig 6 show that POU5F1 and NANOG are activated and suppressed at all times. According to the description of has04550, NANOG and its downstream target genes promote self-renewal and pluripotency. SRF and FOXH1 begin to be activated at 24 h. A full image of the DGRN states is presented in S3 Fig.

Fig 7(b) shows the DGIE scores of the genes in Fig 6. For dataset 3, we focus on hsa04550 signaling pathways regulating the pluripotency of stem cells. Fig 7(b) shows a decreasing tendency, along with Fig 7(a). Once ES cells start to differentiate, the communication ability of the genes in Fig 6 begins to fall. This finding suggests that the activation degree of the regulation of stem cell pluripotency is reduced.

**Fig 7.**
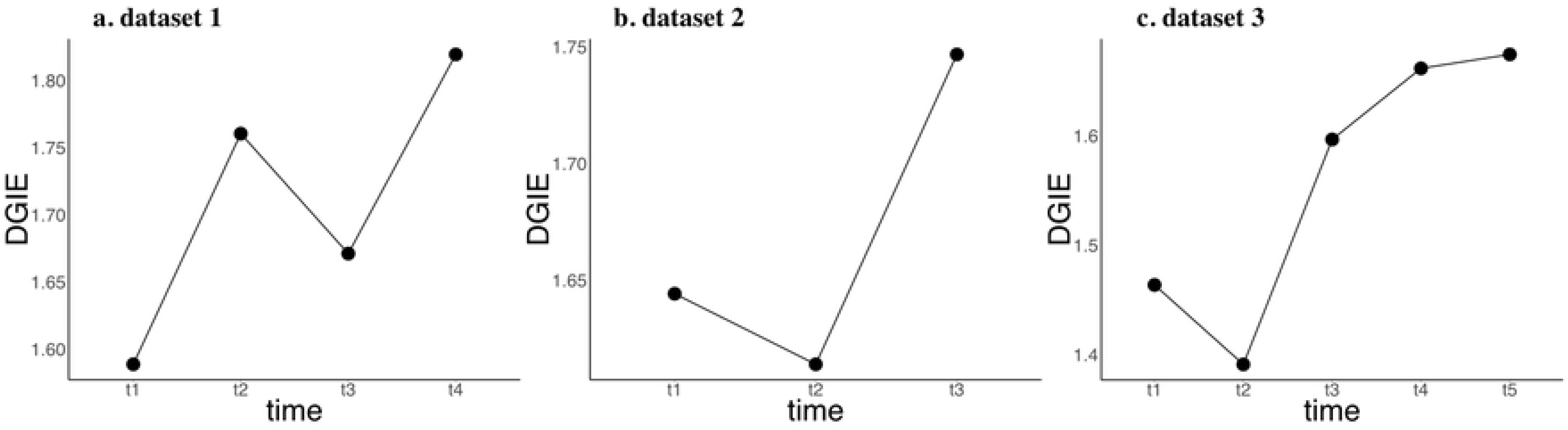
DGIE scores of processes/pathways that are directly related to ES cell differentiation. Datasets 1 and 3 both describe cell differentiation from ES cells. The decreasing tendencies of the DGIE scores indicate that the differentiation functions of ES cells are most active initially and may gradually fade over time.

The DGIE scores of all contributing genes in the DGRN are shown in Fig 8(c). Like datasets 1 and 2, dataset 3 also exhibits an increasing tendency of the DGIE scores. Notably, dataset 3 describes the differentiation of human cells from ES cells. The results help to confirm the conclusions drawn from datasets 1 and 2 with regard to the gradual turn-on of the functions of all known contributing genes.

**Fig 8.**
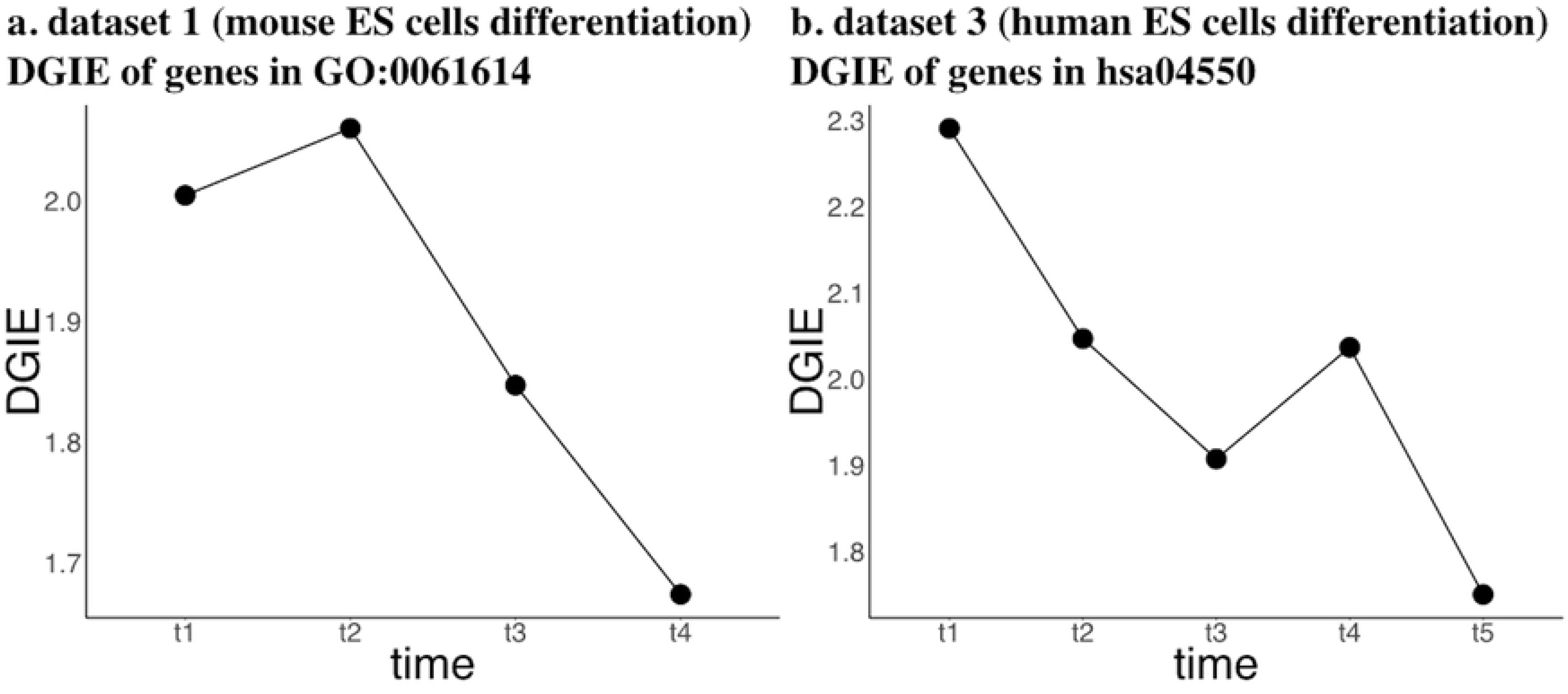
DGIE scores of all known contributing genes. The DGIE scores of all known contributing genes indicate that the communication strength of known contributing genes increases from ES cells to terminally differentiated cells.

S6 Fig shows boxplots of the DGIE scores when a gene or complex is removed from *V*_(1)_. We identify Sox5, Meis2, Hoxb3, Tcf7l1 and Plagl1 as key regulators.

According to the gene annotation information available from the Metascape database, Sox5 is a member of the Sox family, which regulates embryonic development and determines cell fate. Meis2 essentially contributes to developmental processes. Hoxb3 is also involved in development. TCF7L1 plays a role in the regulation of cell cycle genes and cellular senescence. Overexpression of Plagl1 during fetal development causes transient neonatal diabetes mellitus.

The results in S4 Table(C) are similar to those in S4 Table(A), indicating that differentiation-related genes are denser among top-degree genes. S9 Fig shows the differential network of the identified complex, and the counts of the confirmed links in the differential network are shown in S5 Table.

## Discussion

A dynamic network is a powerful tool for elucidating relationships that change over time. With the increasing popularization of single-cell sequencing technology, researchers are obtaining large quantities of time-series single-cell data, which are better able to characterize biological processes than a single snapshot is. To reveal dynamic changes based on time-series scRNA-seq data, we have proposed a novel method of inferring DGRNs with directed links. To ensure that the results are practically and biologically meaningful, we also incorporate the assumptions that the networks are sparse and that consecutive network states are similar into the modeling. Our method, with both the ADMM and PADMM algorithms, shows satisfactory performance on simulated and real datasets.

The greatest obstacle when shifting the level of analysis from bulk data to single-cell-level data lies in the fact that cells are ruined once sequenced by scRNA-seq technology. For this reason, the dynamics at the single-cell level cannot be directly established. Inspired by [15], we first order the cells by their pseudotimes and apply clustering to the ordered cells to obtain groups that can be linked over time. In our algorithm, we specify a number of groups that is equal to the minimum number of cells across all time points in order to use the cell-level information to the greatest possible extent. Because of the complexity of the biological processes, our method may be a simple but compromised approach. The attempt to develop a better way to construct and link cell-level data is an ongoing effort. In practice, when group-level data are available, the proposed method can still be applied by skipping the ordering and clustering steps.

In applications of real time-series scRNA-seq data, it is of interest to characterize changes occurring during biological processes and identify the key regulators. Often, it is difficult to identify these essential differences by inspecting the dynamic graphs themselves (as shown in S1 Fig, S2 Fig, and S3 Fig). The proposed index DGIE serves this purpose by measuring the network differences. In our real data analysis, results obtained based on DGIE scores provide two major insights. First, the DGIE scores of the investigated subnetworks indicate that the differentiation functions of ES cells are most active initially and may gradually fade over time. Second, the DGIE scores of all known contributing genes indicate that the communication strength of known contributing genes increases from ES cells to terminally differentiated cells.

## Conclusion

In this work, we have presented scPADGRN, a novel DGRN inference method using time-series scRNA-seq data. scPADGRN shows advantages in terms of accuracy, robustness and fast convergence when implemented with the PADMM algorithm for network inference using simulated datasets.

In real scRNA-seq data applications, scPADGRN can be used to visualize gene-gene interactions among genes involved in the same biological process or KEGG pathway. These regulation relationships may either persist or disappear.

To quantify network differences, a quantitative index called DGIE has been presented. The DGIE score measures the communication ability of a certain set of genes. At the local level, we have computed the DGIE scores of processes or pathways that are directly related to ES cell differentiation. The decreasing tendency of the DGIE scores indicates that the differentiation functions of ES cells are most active initially and may gradually fade over time. At the global level, the DGIE scores of the three investigated datasets all show the same increasing tendency, indicating that the communication strength of the known contributing genes increases from ES cells to terminally differentiated cells. We have identified a set of genes responsible for changes in the DGIE scores during cell differentiation for each of the three single-cell datasets.

Our results affirm that single-cell analysis based on network inference coupled with quantitative computations can be applied to infer the activity states of gene functions in the process of differentiation from ES cells to terminally differentiated cells, thus potentially revealing key transcriptional regulators involved in cell differentiation and disease development.

In summary, our work provides three main contributions. First, we propose a new method of inferring DGRNs using scRNA-seq data. Second, a quantitative index, DGIE, is proposed to measure the communication ability of a certain set of genes in a DGRN; this index can reflect the activity states of functions in which these genes play a role. Third, key regulators of biological processes can be identified based on the DGIE scores.

## Supporting information

**S1 Fig. Estimated DGRNs for dataset 1**

**S2 Fig. Estimated DGRNs for dataset 2**

**S3 Fig. Estimated DGRNs for dataset 3**

**S4 Fig. Boxplot of DGIE scores after gene/genes removal (dataset 1)**

**S5 Fig. Boxplot of DGIE scores after gene/genes removal (dataset 2)**

**S6 Fig. Boxplot of DGIE scores after gene/genes removal (dataset 3)**

**S7 Fig. Differential network of identified targets for dataset 1**

**S8 Fig. Differential network of identified targets for dataset 2**

**S9 Fig. Differential network of identified targets for dataset 3**

**S1 Table. Full simulation results for Table 1**

**S2 Table. Full simulation results for Table 2**

**S3 Table. Gene lists in dataset 1-3**

**S4 Table. Comparation between rate** 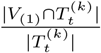, *k* = 10, 50 **and reference rate** 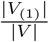

**S5 Table. Number of links and confirmed links in the estimated differential networks.**

## Acknowledgments

This work was supported by the Chinese National Natural Science Foundation (No. 11831015 and No. 61672388) and the National Key Research and Development Program of China (No. 2018YFC1314600).

## References

1. Goode DK, Obier N, Vijayabaskar MS, Liealing M, Lilly AJ, Hannah R, et al. Dynamic gene regulatory networks drive hematopoietic specification and differentiation. Developmental Cell. 2016;36(5):572–587.

2. Wit EC, Abbruzzo A. Inferring slowly-changing dynamic gene-regulatory networks. Bmc Bioinformatics. 2015;16(S6):S5.

3. Li Y, Jin S, Lei L, Pan Z, Zou X. Deciphering deterioration mechanisms of complex diseases based on the construction of dynamic networks and systems analysis. Scientific Reports. 2015;5:9283.

4. Bacher R, Kendziorski C. Design and computational analysis of single-cell RNA-sequencing experiments. Genome Biology. 2016;17(1):63.

5. Shimosato D, Shiki M, Niwa H. Extra-embryonic endoderm cells derived from ES cells induced by GATA Factors acquire the character of XEN cells. Bmc Developmental Biology. 2007;7(1):80.

6. Treutlein B, Qian YL, Camp JG, Mall M, Koh W, Shariati SAM, et al. Dissecting direct reprogramming from fibroblast to neuron using single-cell RNA-seq. Nature. 2016;534(7607):391.

7. Chu LF, Leng N, Zhang J, Hou Z, Mamott D, Vereide DT, et al. Single-cell RNA-seq reveals novel regulators of human embryonic stem cell differentiation to definitive endoderm. Genome Biology. 2016;17(1):173.

8. Matsumoto H, Kiryu H, Furusawa C, Ko MSH, Nikaido I. SCODE: An efficient regulatory network inference algorithm from single-cell RNA-Seq during differentiation. Bioinformatics. 2016;33(15).

9. Cole T, Davide C, Jonna G, Prapti P, Shuqiang L, Michael M, et al. The dynamics and regulators of cell fate decisions are revealed by pseudotemporal ordering of single cells. Nature Biotechnology. 2014;32(4):381–386.

10. Jiao Y, Jin Q, Lu X, Wang W. Preconditioned alternating direction method of multipliers for inverse problems with constraints. Inverse Problems. 2017;33(2):025004.

11. Boyd S CE Parikh N. Distributed Optimization and Statistical Learning via the Alternating Direction Method of Multipliers[J]. Foundations & Trends in Machine Learning. 2010;.

12. Zappia L, Phipson B, Oshlack A. Splatter: simulation of single-cell RNA sequencing data. Genome Biology. 2017;18(1):174.

13. Shannon P, Markiel A, Ozier O, Baliga NS, Wang JT, Ramage D, et al. Cytoscape: A Software Environment for Integrated Models of Biomolecular Interaction Networks. Genome Research. 2003;13(11):2498–2504.

14. Zhou Y, Zhou B, Pache L, Chang M, Khodabakhshi AH, Tanaseichuk O, et al. Metascape provides a biologist-oriented resource for the analysis of systems-level datasets. Nature Communications. 2019;10(1):1523.

15. Duren Z, Chen X, Zamanighomi M, Zeng W, Satpathy AT, Chang HY, et al. Integrative analysis of single-cell genomics data by coupled nonnegative matrix factorizations. Proceedings of the National Academy of Sciences of the United States of America. 2018;115(30):7723–7728.

